# Scarce and directly beneficial reputations support cooperation

**DOI:** 10.1101/788836

**Authors:** Flóra Samu, Szabolcs Számadó, Károly Takács

**Affiliations:** Linköping University, The Institute for Analytical Sociology, 601 74 Norrköping, Sweden; Corvinus University of Budapest, Doctoral School of Sociology, 1018 Fővám tér 8. Budapest, Hungary; Hungarian Academy of Sciences, TK CSS “Lendület’ Research Center for Educational and Network Studies (CSS-RECENS), Budapest, 1097 Tóth Kálmán u. 4, Hungary; MTA Research Group of Ecology and Theoretical Biology, Eötvös Loránd University, Budapest, 1053, Hungary; Budapest University of Technology and Economics, Budapest, 1111, Hungary

**Author notes:** Correspondence and requests for materials should be addressed to F.S.

## Abstract

A human solution to the problem of cooperation is the construction and maintenance of informal reputation hierarchies. Reputational information contributes to cooperation by providing guidelines about previous group-beneficial or free-rider behavior of opponents in social dilemma interactions. How reputation information could be credible, however, when outcomes of interactions are not publicly known, remains a puzzle. In this study, we propose that credibility could be ensured if reputation is a scarce resource and it is not believed to be earned for direct benefits. We tested these propositions in a laboratory experiment in which participants played two-person Prisoner’s Dilemma games without partner selection, could observe some other interactions and could communicate reputational information about possible prospective opponents to each other. We found that scarcity is a necessary condition for reputation scores to be informative. While cooperation has not been sustained at a high level in any of the conditions, reputational information clearly influenced cooperation decisions. The possibility of exchanging third-party information was able to increase the level of cooperation the most if reputation was a scarce resource and contrary to our expectations, when reputational scores have been directly translated into monetary benefits.

## Introduction

Numerous examples demonstrate that social dilemmas are integral parts of our daily lives (Hardin 1968, Axelrod 1984). In a social dilemma, there is a conflict between individual and community interests; for instance, when citizens consider their participation in social movements (Olson 1965) or in the case of the overconsumption of common resources (Hardin 1968). The most severe cases of these kind of conflicts in which following selfish interests is the dominant strategy that leads to an outcome that is disadvantageous for everyone in the group have been modelled as the multiplayer Public-Goods Game (PGG) or the Prisoner’s Dilemma (PD) game (Luce and Raiffa 1957). Over the past decades, dozens of researchers have been attracted by the problem leading to a wide range of suggestions for solutions to social dilemmas (Nowak 2006, van Lange et al. 2014).

One of the proposed informal solutions is the establishment and maintenance of reputations that provide guidelines for appropriate actions towards interaction partners. In large populations it is difficult to observe the past decisions of the potential unknown transaction partners directly. Under such conditions, cooperation can be established through the use of reputations that trigger conditional cooperative behavior (Wedekind and Milinski 2000, Milinski et al. 2002, Panchanathan and Boyd 2004, Fu et al. 2008, Feinberg et al. 2014). Individuals strive for reputations as they would like to harvest the benefits of cooperative interactions. In turn, they can earn their reputations by cooperating with others. Empirical studies confirmed that people increase their cooperative behavior when they are observed by others (Semmann et al. 2004, Burton-Chellew, El Mouden and West 2017) and even in the presence of a depicted eye (Haley and Fessler 2005, Mifune, Hashimoto and Yamagishi 2010).

Why would individuals display cooperative behavior publicly? The reputation they gain might help them in the process of partner selection (Bliege Bird and Powel 2015), acquiring resources (Willer 2009), gaining social network benefits (Lyle and Smith 2014), greater influence (Milinski, Semmann, and Krambeck 2002), or accumulating other rewards (Frey and van de Rijt 2016). A further possible answer is that reputation they earned by their pro-social behavior helps them to receive pro-social acts from others: ‘I scratch your back and someone else will scratch mine’ (Nowak and Sigmund 2005: 1291). These future expected returns could rationalize cooperation already in the first interaction. Cooperation at the outset could work as a signal of future commitment (Wedekind and Braithwaite 2002, Bliege Bird and Smith 2015) or of risk taking for the common benefit (Willer 2009). Either way, it is a costly investment that reflects some form of trustworthiness (Molm 2000).

In previous models and experiments, where reputations have been shown to provide an efficient solution for social dilemmas, individuals could observe the past behavior of others and hence they had perfect and complete information on who has been cooperating and who has not (Wedekind and Milinski 2000, Milinski et al. 2001, Milinski, Semmann and Krambeck 2002, Semmann, Krambeck and Milinski 2005, Seinen and Schram 2006). In such circumstances, the reputation of an individual can simply be expressed in an image score that conveys true information about past behavior (Nowak and Sigmund 1998, Milinski et al. 2001, Nax et al. 2015). In practice, however, the outcome of all transactions or a credible summary score is not publicly available. The mechanism that helps to access reputational information is informal communication in which individuals exchange their own previous experience and the information they have received with others. Such an evaluative exchange of third-party information is referred to as gossip (Foster 2004). In experiments where participants could gossip to each other in the form of sending messages, transmitted information was very much in line with observed choices (Sommerfeld, Krambeck, Semmann and Milinski 2008, Feinberg 2014). Moreover, gossip was credited even when it was unverifiable (Fehr and Sutter 2019), and when direct observation was also possible (Sommerfeld et al. 2007).

In line with the literature that recognizes the importance of gossip and reputation for cooperation, we expect that where reputation building is available, individuals endeavor to increase or maintain their reputation by cooperation. The assumption of strategic reputation building claims that individuals intend to increase their own reputation to make transactional partners more willing to cooperate with them.

### H1. The possibility of gossip increases cooperation compared to the baseline rounds without communication

In this study we are able to test how gossip dynamics works. On the one hand, individuals use gossip as an informal institution, and hence they transmit information about others’ cooperative behavior through gossip. Thus, we expect that gossip will be in line with previous decisions of the partners.

### H1a Individuals use gossip to transmit their direct observations to others

On the other hand, receivers of gossip have to rely on the information they received. We therefore expect that they adjust their private reputational evaluations about others in line with the gossip received.

### H1b Received gossip alters individuals’ private reputational evaluation on others

Information received through gossip supplements direct observations and allows for a better alignment of decisions in the social dilemma towards the interaction partner. When interactions are not observed publicly, however, there is no guarantee that the received information is credible and truly reflects merits from previous actions. The possibility of misrepresentation and strategic deception undermine the sustainability of cooperation through reputation (Számadó, Szalai, and Scheuring, 2016). How reputational information could be credible in games of conflict of interest is still an unanswered question in the literature.

In this study, we propose that credibility of the reputational system could be ensured if reputation is a scarce resource. The link between the scarcity of reputations and cooperation is not evident just as the effect of competition induced by the scarcity and cooperation. On the one hand, when reputations are expressed in relative terms, they better differentiate between the individuals and hence create information content for their value. As they are able to differentiate more between partners than reputation scores that are infinitely available, they become more reliable sources of information. Hence, individuals compete for being more cooperative than others in order to keep up with their reputation (Barclay 2013). The relativized interpretation of reputations is more in line with the everyday use of the reputation concept (Paine 1967). On the other hand, when reputations are scarce, there is more motivation for strategic behavior for acquiring them. This includes a rational justification of the strategic undermining the reputation of others.

The theory of ‘competitive altruism’ (Roberts 1998, Barclay 2004, Barclay 2013, Roberts 2015, Simpson and Willer 2015) states that expectations of indirectly reciprocated help doesn’t explain satisfactorily the motivation to pro-social behaviors – as the theory of indirect reciprocity claims it is – but rather the competition is the motivation for reputation-building which incorporates in the rise of pro-social behavior (Feinberg et al. 2014). The scarcity of reputational resources creates competition between individuals where higher reputational position requires more strategic investment, which can be achieved with a higher level of cooperation. Thus, we expect that in a competitive environment for reputation cooperation will increase.

### H2 Competition for scarce reputations increases cooperation

As another mechanism, we propose that reputations are reciprocated with cooperation once they are believed that have not been earned for direct benefits, but they are the result of voluntary contributions to the collective good (Smith and Bliege Bird 2000, Bliege Bird and Power 2015, Raihani and Bshary 2015). Empirical evidence finds that pure reputational rewards that have little chance to materialize in the future (e.g. prizes without monetary benefits) still can facilitate pro-social acts (Gallus 2016). Considering one type of reward-based investment schemes, customer loyalty programs shows that promoting relationship-oriented rewards – compared to direct monetary rewards – attracts more investment (Roehm et al. 2002). Moreover, which is even more interesting, the most successful crowdfunding projects allow self-selection into relatively small, distinct types of reward tiers: lower investments with tangible benefit or higher investment with intangible rewards (Shi 2018).

External incentives have an ambiguous effect on cooperation. On the one hand, tangible incentives can foster cooperation simply because they reduce the magnitude of conflict between self-interest and the common good (Rapoport and Chammah 1965). On the other hand, empirical studies have shown that external incentives can reduce the motivation for reputation-building (Frey and Jegen 2001, Bowles 2008, Bravo, Squazzoni, and Takács 2015) and as a result, the level of cooperation does not grow as much as we would observe in the absence of this ‘crowding-out effect’ (Bowles and Polanía-Reyes 2012). Either punishment as an external incentive (e.g. Deci et al 1999, Mulder et al., 2006) or rewards (e.g. Ariely, Bracha, and Meier 2009) can reduce motivation to achieve high reputation. The mechanism behind the reduced motivation for reputation-building supposed to be the lack of opportunity to signaling group-based motivation or commitment in the future (Johansen and Kvaloy 2016, Gneezy 2011).

We expect that if direct external incentives are linked to the reputational position, then the signal of long-term commitment or group-based motivation will be inseparable from the motives for direct benefits. In this case reputational signals will be less efficient.

### H3 Direct monetary stakes for reputation decrease cooperation

We aim to show how extrinsic motivation and competition for scarce resources affect strategic reputation building and cooperation in an environment where there is a low probability of meeting with the same person again. We tested whether the degree of competition influences the level of cooperation by alternating two elements of a reputation system, namely the scarcity of the reputational resources (H2) and further external incentives for reputation-building (H3). We expect that the highest cooperation level will appear under the condition when individuals are managing scarce resources, while the lowest will happen in a monetarily incentivized context where partners’ evaluations are not relative and therefore competition is less intense.

## Methods

### Participants

We investigated our hypotheses in anonymous two-person Prisoner’s Dilemma games in an experimental laboratory with volunteer participants. The experiment was conducted at the Corvinus University of Budapest between 13-25 November 2016. In total, 160 individuals participated in the experiment (male: 54.4%) in eight sessions in groups of 20.

### Design

The experiment followed a 2×2 between-subject design. The scarcity of reputations was manipulated by the way how participants could distribute reputation scores to others (on a scale between 0 and 100). Participants could either allocate a fixed amount of reputation scores (*scarcity*) or there was no ceiling on the distributable scores (*abundance*). Direct benefits for reputation was manipulated as reputation scores were either symbolic (*not paid*) or were incentivized financially (*paid well*). In the latter case, participants received the payoffs from the PD games and nothing more or less if they received 50 reputation points (the midscale value) from other participants on average. Otherwise, a one-unit decrease/increase from the default value of 50, reduced/increased their payments by HUF 20 (∼ EUR 0.06). For instance, if all participants gave zero reputation to someone, then the receiver’s payment has been decreased by HUF 1000 (∼ EUR 3.12). The four experimental groups are the combination of these two manipulations.

### Procedure

The experiment has been programmed using the experimental software z-Tree (Fischbacher 2007). Participants have read the instructions on paper and on their screen after they have been randomly assigned to a computer in the lab. Subsequently, they had to fill in a quiz of understanding and when in doubt, could ask questions privately.

In the experiment, participants were identified with numbers ranging from 1 to 20. In each round, they were randomly shuffled and have been matched with two other participants in each round whose IDs were displayed on participants’ screen. Participants played the two-person Prisoner’s Dilemma (PD) with the two opponents parallel. PD options were labeled with ‘L’ and ‘R’. The cooperative decision was marked with ‘L’. PD payoffs through the experiment were fixed to HUF 1500 (∼ EUR 4.7) for mutual cooperation; HUF 500 (∼ EUR 1.6) for mutual defection; HUF 2500 (∼ EUR 7.8) for temptation; and 0 otherwise. They had 23 seconds to decide. If they ran out of time, they received nothing, hence time-out could not be considered as an opt-out or walk away strategy (Hauert et al. 2002, Aktipis 2004, Rand and Nowak 2011). Results appeared on the screen after every PD game.

In the first five baseline rounds, participants played the PD games without communication. From the 6^th^ round on, they played the PD game for several rounds with gossip in one of the four experimental conditions. In each round, in the presence of reputation scores for others on their screen they played the two two-person Prisoner’s Dilemma (PD) games simultaneously (1^st^ screen). Then, they could observe the decision of two players as an outside third party (2^nd^ screen). Hence, in all four treatments participants were able to observe the PD decisions of four players in total: their two interaction partners and two randomly selected players. Subsequently, they could send gossip to a randomly selected participant, whose ID has appeared on their screen (3^rd^ screen). In four empty boxes they could enter the ID of those targets they wanted to send a message about. Under the boxes, participants could select positive, neutral, negative emoticons as the gossip message. Sending a gossip was optional, and it was free of charge. The gossip messages were received, and participants could assign reputation scores to all participants subsequently (4^th^ screen). The instructions for reputation scores asked participants to evaluate how trustable they think others are on a scale of 0 to 100. At the end of each round, the average of their received reputation points was displayed (5^th^ screen).

The final profit was calculated as an average payoff of 6 randomly selected rounds. In addition to the final payoff, a show-up fee (HUF 1000) has been paid to the participants. Participants earned 1822 HUF on average.

## Results

### Cooperation

Figure 2 depicts the average cooperation level across time groups in the four treatment groups. At first sight, we see higher cooperation level in Round 6 (M=26.8, SD=44.4, incl: M=29.9, SD=45.8) compared to the first five rounds where communication was not allowed (M=22, SD=41.5, incl: M=24.0, SD=42.7). Inspecting how decisive this result is, we run multilevel logistic regression analysis which revealed that the possibility of information exchange increased cooperation significantly only in the ‘competition-paid well’ treatment (see Table A1a and A1b in Appendix 1), which means only a partial confirmation of Hypothesis 1.

**Figure 1.**
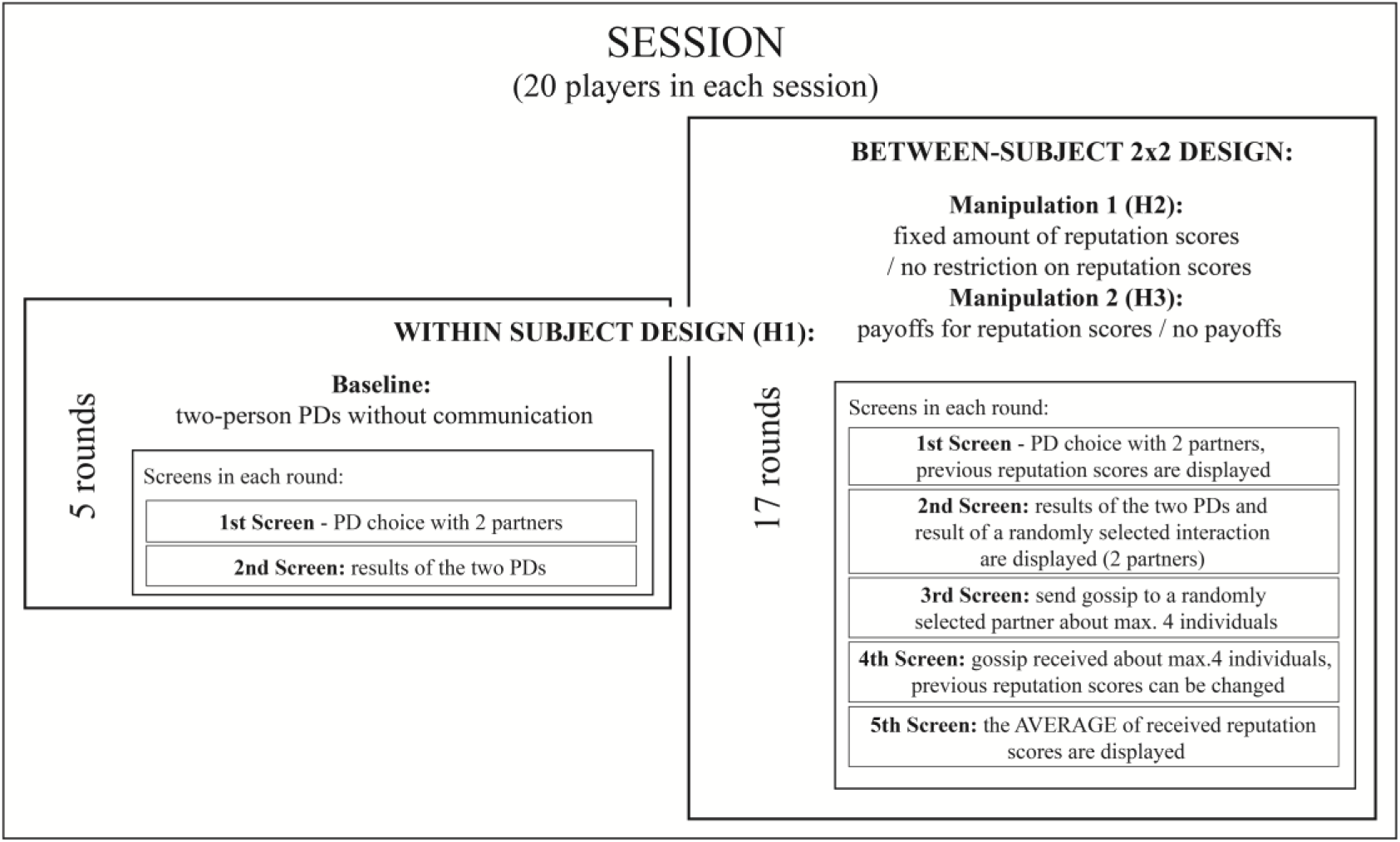
Design of the experiment through an example of one session.

**Figure 2.**
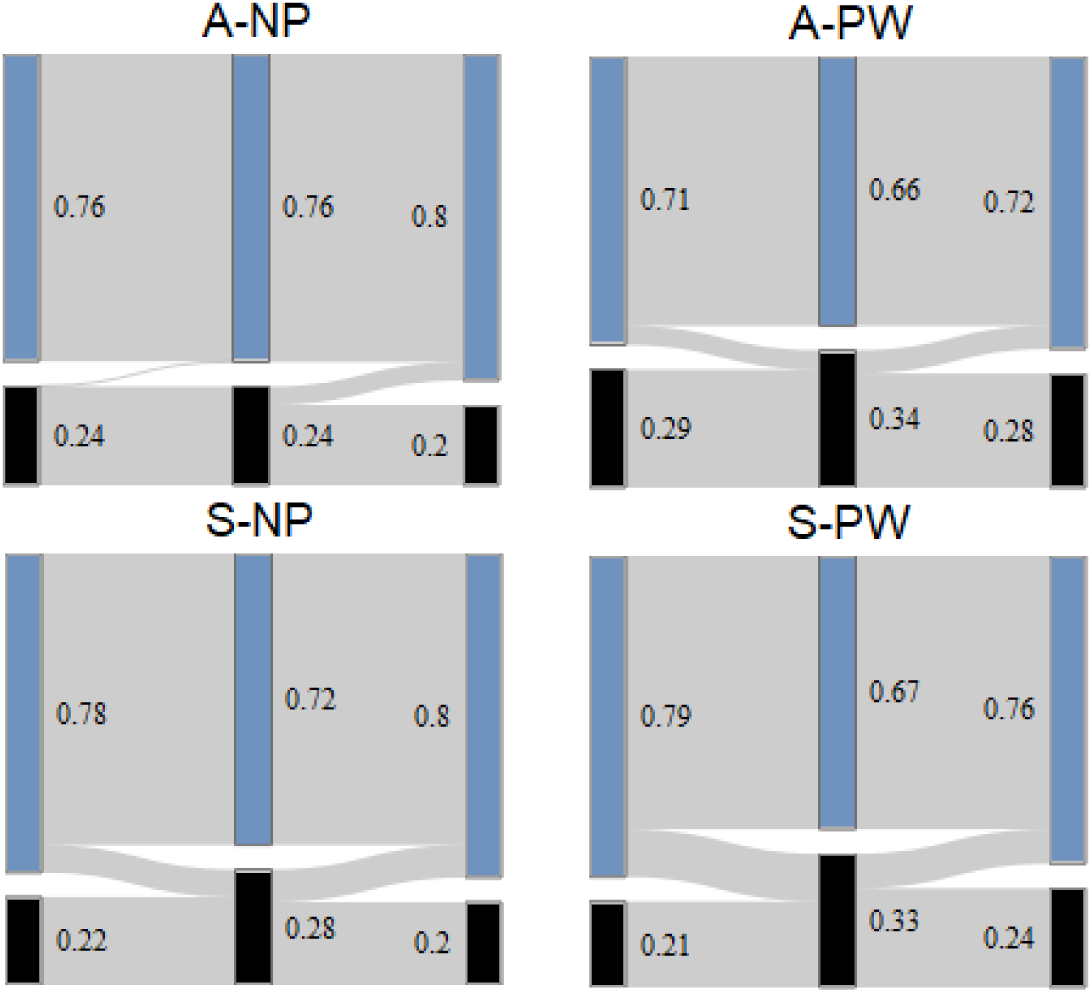
Cooperation level in treatments.

As regards Hypotheses 2 and 3, neither the implementation of competition nor the introduction of incentives was sufficient in itself to increase cooperation in the group. The two manipulation together, however, was able mitigate a declining tendency of cooperation over time (Van Lange 1999) compared to what we observe in the ‘abundance – not paid’ treatment (see treatment – round interaction effect in Table 1b in Appendix 1). This means that if reputational system contains both competition and external incentives (*‘scarcity – paid well’*) cooperation survive longer than without these features (*‘abundance – not paid’*). The presence of only one of these elements ranks the two other treatments between the two poles based on the observed level of cooperation. In the light of these results, it appears that scarce reputations with direct incentives together could help to maintain cooperation, however, this condition also failed to increase the level of cooperation in the long run. These results suggest that reputations could be more credible and efficient pathways to cooperation if they are relativized and create competitive dependences between participants. Why we do not see a larger impact is potentially due to the fact that competitive dependencies encourage participants to send dishonest reputational information about others and the higher role of strategic misrepresentations could undermine the credibility of information and hence reputation-based cooperation.

**Table 1.**
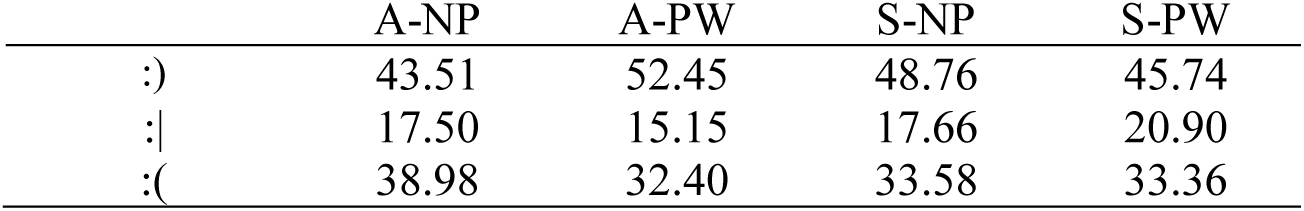
The distribution of gossip values by treatments

A reputational system is reliable if it appropriately reflects the potential behavior of others that is otherwise hidden to new partners. It functions well if it helps individuals to cooperate with those who have a higher reputation and defect against those who have lower willingness to cooperate. Figure 3b depicts the average degree of cooperation with alter as a function of alter’s cooperation level during the whole experiment. Since there was no signaling system in the baseline rounds, it is not surprising that in the first five rounds in each treatment there is no correlation between someone’s cooperative behavior and the partner’s average behavior (corr = −0.07, *p* = 0.38). We found small, but significant correlations in two treatment groups: in the ‘*abundance – not paid*’ (corr= 0.31, *p*= 0.05) and in the ‘*scarcity – paid well*’ (corr= 0.32, *p*= 0.04) treatment (see more detailed analysis Table A2 in Appendix I). These correlations illustrate how well the reputational system have reflected others’ previous behavior, which is a prerequisite for the operation of reputation systems. The correlations suggest that the reputation system has improved its credibility within the time available.

**Figure 3.**
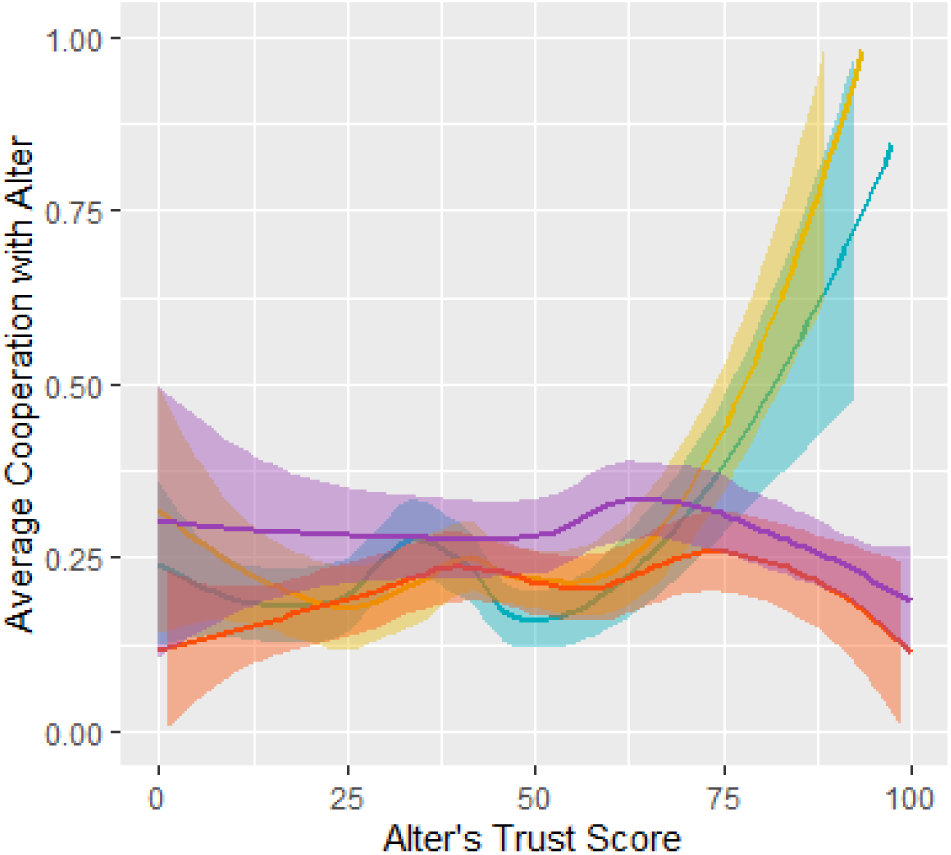
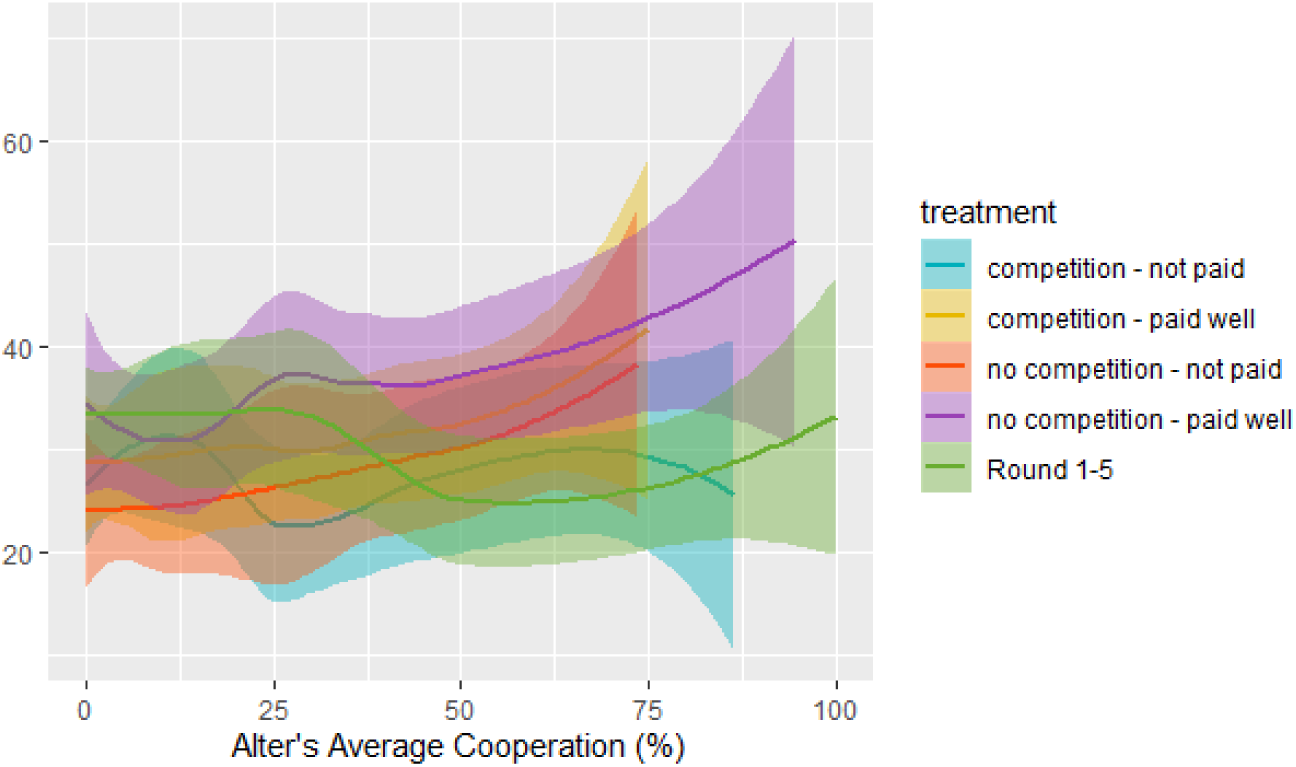
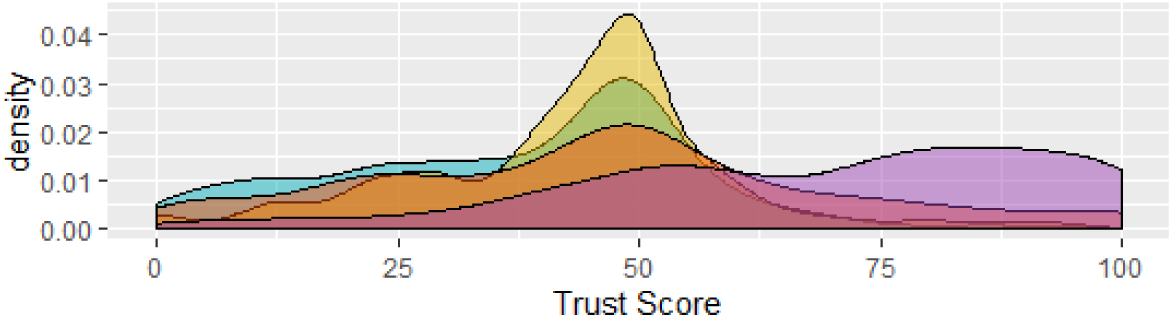
The accuracy of the reputation systems by treatments.

Examining the consistency of reputational rankings among participants – to see how far they have reached consensus in ranking participants in their private reputational scale – we calculated the inconsistency of participants’ rank between others’ reputational hierarchy in each treatment:

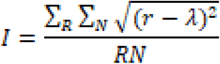

where *λ* is the average rank of a given individual in all reputational hierarchy in a given round, r is the rank of the same individual in subjects’ reputational scale in the same round, N is the number of subject and *R* is the number of rounds. A t-test performed between each treatment showed no significant differences (p>0.05) (see Figure A1 in Appendix 1).

But we see significant differences in how the distribution of reputation scores evolved over time by calculating the stability of reputation scores received between rounds:

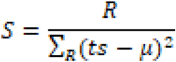

where *S* is the stability of objects’ reputation score, *R* is the number of rounds, *µ* is objects’ average reputation score throughout the entire experiment and *ts* is the score that others gave to others in one round. The smallest variation can be found in the ‘*scarcity – paid well*’ treatment (M= 0.65, SD=0.21), which is significantly different from the ‘*scarcity – not paid*’ treatment which has the second highest stability (M= 0.62, SD=0.21, t-tests p<0.001). The *‘abundance – not paid’* treatment is the third in the stability ranking which also differs significantly from all other treatments (M=0.61, SD= 0.20, t-tests p=0.03). The *‘abundance – paid well*’ has significantly less consistency on the other side of the scale, where there was no restriction on how many reputational score a participant can distribute among other participants (M=0.56, SD= 0.21, t-tests p<0.001). In the following section we examine further how direct observations of participants and indirect information received from others contributed to the formation of the reputational hierarchy (see Figure A2 in Appendix 1).

### Reputation

The average reputation score players gave to each other is slightly lower than the initial value of 50 (M = 46.1, SD = 29.9), and the way how reputation scores distributed among participants vary in the different treatments (see Figure 3). Treatment groups without competition (*‘abundance’* treatments) have been characterized by higher scores than the initial value, since in these treatments, participants could give high scores without lowering the scores of other players. The presence of lower points was more typical in the *‘not paid’* treatments, where scores did not affect participants’ payoffs of the PD game. These trends observed along the two manipulations causes the difference in the distribution of reputation scores between the four treatments: the ‘*abundance–paid well’* treatment was characterized by merely high scores (M=67.8% SD=35.1) and the ‘*scarcity - not paid*’ group by rather low scores (M=37.4% SD=24.6). In sessions where reputation scores could be distributed infinitely and players were *not paid* after their average reputation score we see a wide-spread scoring in both low and high directions (M=45.7% SD=31.2), while in the ‘*scarcity – paid well*’ treatment we observe a less extreme negative shift in the reputation score distribution (M=43,0% SD=23.5).

We estimated a multilevel linear regression model to explain the allocation of reputation scores. It is natural that participants based their scores on previous values. We found that the cooperative decisions of interaction partners and observed players, as well as positive messages had a positive effect on the allocated reputation scores, while defections and negative messages negatively influenced scores (see Model A2, Table A5, Appendix I). A higher number of messages sent by gossipmongers has not been rewarded with higher reputation scores (see Model A2, Table A5, Appendix I).

Looking for differences between the treatments we found that in the ‘*abundance-paid well*’ condition negative gossip generated a greater volume of score reduction in comparison to other treatments and neutral gossip have also negatively affected the points in this group in comparison to other treatment groups. Positive gossip was the most powerful in the scarcity-paid well treatment. Competition for scare reputation scores together with incentives reduce the reliance on negative gossip, probably because in this treatment players are encouraged to decrease the reputation of others to improve their own position, by sending negative gossip about their rivals.

### Gossip

In the experiment, participants could share reputational information or strategically misrepresent others only in a reduced form. They could send messages of three types about the selected target: a happy, a neutral, or a sad smiley. 47.7% of the possible messages were sent by the participants. (A-NP: 47.5%, A-PW: 44.3%, S-NP: 53.9%, S-PW: 45,2%) Participants had one minute to send messages. In the very beginning many players run out of time, but this has improved soon and on average only 15% of players used all the available time to send out messages, which could mean that lack of time was not the reason for the sparsity of gossiping.

A multilevel ordered logistic regression model (see Table A6, Appendix I) revealed the same pattern in gossiping what we can observe in Figure 3. A one unit increase in previous reputational points increased the probability of a positive and reduced the probability of negative messages (see Table A6, Appendix I). This positive correlation remains significant in all treatment groups after controlling for the observed outcomes of PD games. In addition, gossip ratings were very much in line with observed choices (Sommerfeld et al. 2007): a positive message was more likely to be sent if someone was cooperative, and a negative message if someone did not cooperate. We have found no evidence that cooperative individuals are more likely to be the target of the gossip (antisocial punishment). The higher evaluation of individuals with low level of reputation in competition could arise by the fact that more messages were sent randomly in the scarcity treatments probably because the competitive situation has been perceived as more complex.

At the end of the experiment, participants were asked how motivated their choices were. On a seven-point scale between the statements of ‘I always sent messages randomly’ and ‘I always sent messages with a purpose’ the mean value is towards consciousness in every treatment (M = 4.95, SD = 1.98). The observed L or R decisions - also measured on a 7-degree scale - played only a minor role in sending messages (M = 4.8, SD = 2.2). In response to an open question about the reasons of sending messages, 15.7% mentioned that they sent random messages. 46.4% of the participants felt that they were unable to influence others with their messages.

## Discussion

In this study we attempted to investigate how cooperation can be sustained by private reputations. Once public information is not available about previous actions, reputations could guide conditional cooperation towards interaction partners. In line with our expectations (H1) we found an increase in the level of cooperation when the institution of reputations and gossip have been introduced, though in most treatments, cooperation has faded over time, which is a typical feature of Social Dilemma experiments (Fischbacher et al. 2001). Reputations, however, could be distorted or strategically influenced. We proposed two mechanisms that could safeguard the credibility of reputational information. First, we investigated whether the scarcity of reputational resources (H2) with the expectation that reduced access will increase competition, could increase reputation-based cooperation. Second, we analyzed whether additional monetary incentives connected to reputation would distort the credibility of reputations (H3). In our experiments, we have not found a support at all for the latter hypothesis (H3). Instead, we found that at the intersections of these two manipulations, competition for scarce monetary rewards has resulted in higher cooperation that was sustained at a relatively high level in the long run.

If we have a closer look at the setting with scarce reputational resources that could directly be translated to monetary gains, then we can envisage a fierce competition in which rational individuals would compete to maintain or increase their payoffs by either increasing their own cooperative behavior or by wrecking the position of others. Therefore, when the rewards and punishment are intense, we expected that cooperation will be higher, and competitors will use the opportunity to worsen others’ position to avoid material injury. We found some evidence of this as we observed more cooperation over time than in other conditions and we also found a slightly different pattern in participants’ reliance on gossiping. It is important to note that the dissemination of false information cannot just contribute to the deterioration of reputation, it can also hinder the reliability of the reputation system. Since individuals are only willing to take the risk of cooperative behavior if the reputation system is reliable and if others rely on it as well, this can have a negative effect on cooperation. Hence, the reputation system could be less reliable and thus cooperation might collapse. The collapse of the reputation system, however, did not happen, in our experiment. The evidence suggests precisely the opposite, among all treatments the level of cooperation here has the smallest decrease over time. It seems that in this treatment, participants took the competition more seriously and strived more for achieving their own reputations by cooperation rather than by downgrading the reputation of others, possibly by realizing the riskiness of the latter route. Reputations have preserved their credibility over time despite the possibility of misinformation as negative gossip was not always believed and have not always been taken into account at decisions.

We found that the use of external incentives without competition is hampering the development of a trusted reputation system. Under such circumstances, individuals did not rely on reputation scores during their PD decisions. On the one hand, with unlimited reputational resources, individuals tried to encourage cooperation, thus they gave high reputation points for everyone, making it impossible to use the reputational system to differentiate cooperative intentions of others. In the condition with abundant reputation scores with direct monetary rewards, one possible explanation for not finding a link here can be the generosity of scoring. In this treatment the allocation of reputation scores was not just unlimited, but participants could earn extra money based on their average scores, therefore participants might have tried to incentivize others and gave almost the maximum for everyone. By doing so, they might have lost a benchmark which helped them to decide whom to cooperate with.

The reputation system was unable to reflect cooperative behavior where limited reputational resources were available, but there was no external motivation for reputation. Here the problem probably was that there was no need to strive for the scarce reputational scores, and competition has not been taken seriously by participants.

In general, the fact that we do not experience a larger impact of reputations on cooperation in our experiment could be attributed to several reasons. Primarily, we investigated the two-person PD game with random reshuffling of partners and no publicly available information, which itself is the most severe social dilemma in which rational action is simply defection. It is possible that the magnitude of conflict in the Prisoner’s Dilemma game was so strong that even a well-functioning reputation system couldn’t increase cooperative acts. Unfortunately, in this study it is impossible to assess whether the magnitude of the conflict is responsible for the low level of cooperation. It is also possible that reputation scores worked to a limited extent because of the abstract situation. The credibility of reputations has probably also been prone to error in our experiment. Potentially, participants had problems to remember their earlier experiences and might have also mixed up other participants as they were identified with numbers that are harder to recall than names or faces. It is also difficult to assess whether participants used second-order norms (Ohtsuki and Iwasa 2004, 2006, Számadó, Szalai, and Scheuring 2016) that have also lowered cooperation - and in some of the experimental conditions more than in others. For instance, it is possible that participants punished individuals who cooperated against participants with a low reputation score with defection and they did this most frequently when reputations were scarce.

Still, our results bring us closer to understanding under which conditions reputations and gossip contribute to cooperation. Further research is needed to find out, however, under which conditions gossip is used strategically and in a dishonest way to undermine the reputation of others, and under which conditions it could be considered as altruistic punishment of defectors (Feinberg, Cheng, and Willer 2012, Feinberg, Willer, and Schultz 2014).

## Acknowledgements

The project has received funding from the European Research Council (ERC) under the European Union’s Horizon 2020 research and innovation programme (grant agreement No. 648693) The authors gratefully acknowledge support from the Hungarian National Research, Development and Innovation Office NKFIH (grant number K112929). We would like to acknowledge and thank Béla Janky and Róbert Tardos for their assistance in the interpretation of the results and students at the Corvinus University Budapest for their participation in the experiment.

## Author contributions statement

K.T. designed research; F.S. performed and analysed the results; F.S., K.T. and Sz.Sz. wrote the paper.

## Appendix I

**Table A1a.** Estimated coefficients with multilevel mixed-effects logistic regressions (separate models for treatments)

Appendix I. **Table A1a Estimated coefficients with multilevel mixed-effects logistic regressions (separate models for treatments)**
| Dependent variable: Prisoner's Dilemma choice<br>0 = defect, 1 = cooperate |  | abundance –<br>not paid | abundance –<br>paid well | scarcity –<br>not paid | scarcity –<br>paid well |
| --- | --- | --- | --- | --- | --- |
| baseline: Round 1-5 |  |  |  |  |  |
|  | Round 6 | 0.16 | -0.30 | 0.45 | 0.91* |
|  | Round 7 | -0.43 | -0.05 | 0.29 | 1.16*** |
|  | Round 8 | -0.65 | 0.19 | 0.40 | 0.27 |

**Table A2b.** Estimated coefficients with multilevel mixed-effects logistic regressions.

| Dependent variable: Prisoner's Dilemma choice<br>0 = defect, 1 = cooperate | Model 1 |
| --- | --- |
| baseline: abundance - not paid |  |
| abundance - paid well | -0.11 |
| scarcity - not paid | -0.45 |
| scarcity - paid well | -0.32 |
| baseline: Round 1-5 |  |
| Round 6 | 0.16 |
| Round 7 | -0.28 |
| Round 8 | -0.52 |
| Round 9-23 | -0.45** |
| treatment – round interaction |  |
| abundance - paid well # Round 6 | -0.39 |
| abundance - paid well # Round 7 | 0.25 |
| abundance - paid well # Round 8 | 0.76 |
| abundance - paid well # Round 9-23 | 0.47 |
| scarcity - not paid # Round 6 | 0.25 |
| scarcity - not paid # Round 7 | 0.56 |
| scarcity - not paid # Round 8 | 0.90 |
| scarcity - not paid # Round 9-23 | 0.19 |
| scarcity - paid well # Round 6 | 0.71 |
| scarcity - paid well # Round 7 | 1.38** |
| scarcity - paid well # Round 8 | 0.79 |
| scarcity - paid well # Round 9-23 | 0.61*** |
| _cons | -1.71*** |
| lns1_1_1 | 0.59*** |
| Nr. of groups | 160 |
| N of obs | 6326 |

**Table A3.** Estimated coefficients with multilevel mixed-effects logistic regression.

| Dependent variable: Prisoner's Dilemma choice<br>0 = defect, 1 = cooperate | abundance –<br>not paid | abundance –<br>paid well | scarcity –<br>not paid | scarcity –<br>paid well |
| --- | --- | --- | --- | --- |
| Alter's cooperative behavior in previous rounds | 1.15*** | -0.06 | 0.26 | 0.85* |

**Table A4.** Multilevel mixed-effects linear regression.

| **Alter's cooperative behavior in previous rounds (%)** | **Model 1** |
| --- | --- |
| baseline: abundance - not paid |  |
| abundance - paid well | 0.05*** |
| scarcity - not paid | -0.02*** |
| scarcity - paid well | -0.02** |
| Alter's Reputation score | 0.00*** |
| Interaction |  |
| abundance - paid well * Alter's Reputation score | -0.00 |
| scarcity - not paid * Alter's Reputation score | 0.00*** |
| scarcity - paid well * Alter's Reputation score | 0.00*** |
| _cons | 1.20*** |
| lns1_1_1 | -5.55*** |
| lnsig_e | -1.42*** |
| N | 48758 |

**Table A5.** Multilevel mixed-effects logistic regression.

| **Dependent variable: Prisoner's**<br>**Dilemma choice**<br>**0 = defect, 1 = cooperate** | **Model 1** | **Model 2** | **Model 3** | **Model 4** | **Model 5** |
| --- | --- | --- | --- | --- | --- |
| baseline: abundance - not paid |  |  |  |  |  |
| abundance - paid well | 0.38 | 0.41 | -0.32 | 1.82* | 1.83* |
| scarcity - not paid | -0.18 | -0.17 | 0.06 | 0.55 | 0.55 |
| scarcity - paid well | 0.46 | 0.45 | 0.58 | 0.55 | 0.49 |
| Alter's cooperative behavior in previous rounds |  | 0.68*** | 0.30 | 0.25 | 0.19 |
| Alter's Reputation score |  |  | 0.03*** | 0.04*** | 0.04*** |
| abundance - paid well * Alter's Reputation score |  |  |  | -0.03*** | -0.03*** |
| scarcity - not paid * Alter's Reputation score |  |  |  | -0.01 | -0.01 |
| scarcity - paid well * Alter's Reputation score |  |  |  | 0.00 | 0.00 |
| baseline: Partners defect in both PD game (in the previous round) |  |  |  |  |  |
| One partner cooperates, the other partner defects |  |  |  |  | 0.26* |
| Partners cooperate in both PD game |  |  |  |  | 0.49* |
| baseline: Not observed |  |  |  |  |  |
| Observed participant defect |  |  |  |  | -0.25 |
| Observed participant cooperate |  |  |  |  | 0.43 |
| Round |  |  |  |  | -0.02* |
| _cons | -2.22*** | -3.07*** | -4.14*** | -4.52*** | -4.20*** |
| lns1_1_1 | 0.66*** | 0.66*** | 0.77*** | 0.78*** | 0.78*** |
| N | 4326 | 4326 | 4326 | 4326 | 4318 |

**Table A6.**
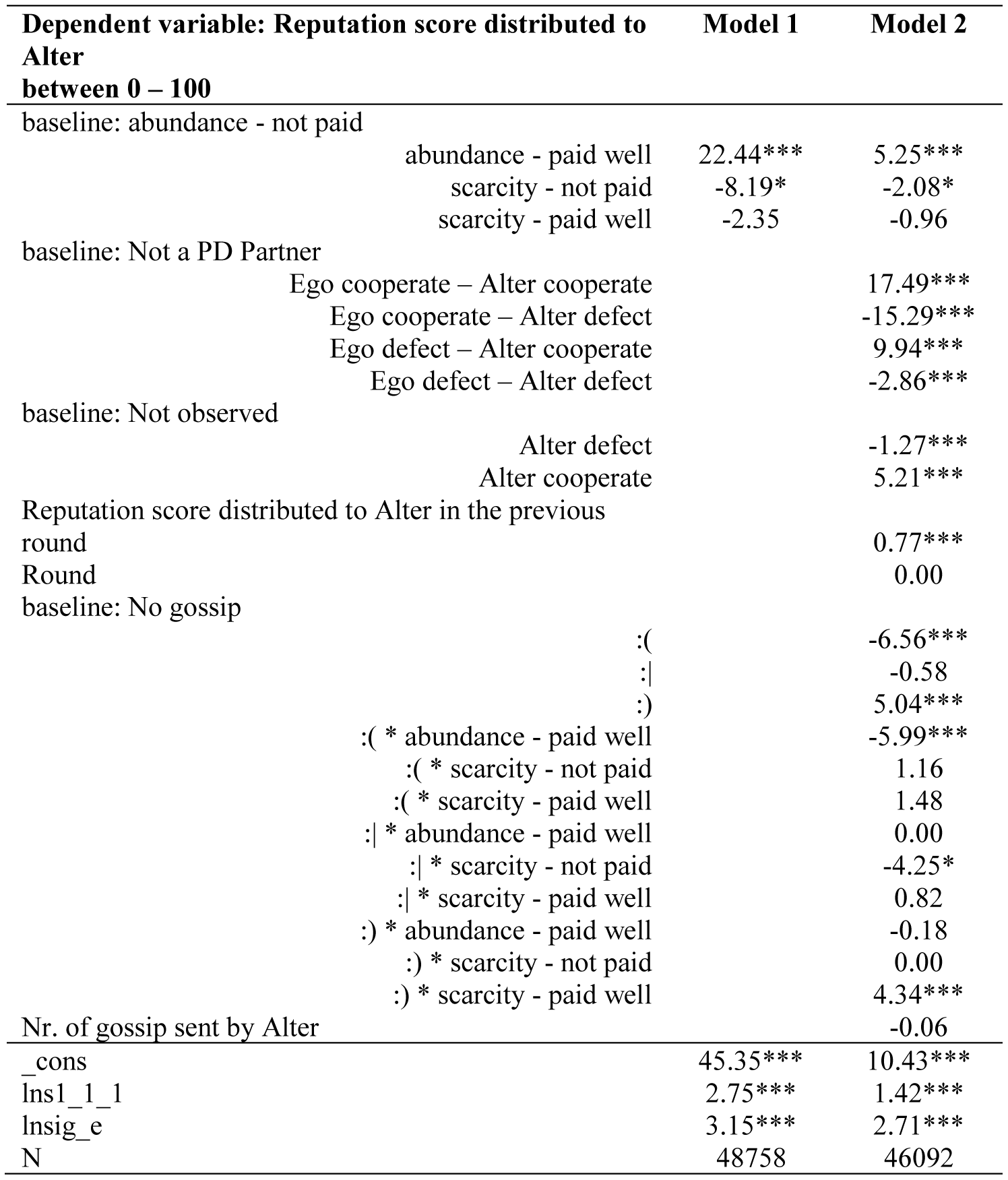
Multilevel mixed-effects linear regression.

**Table A7.**
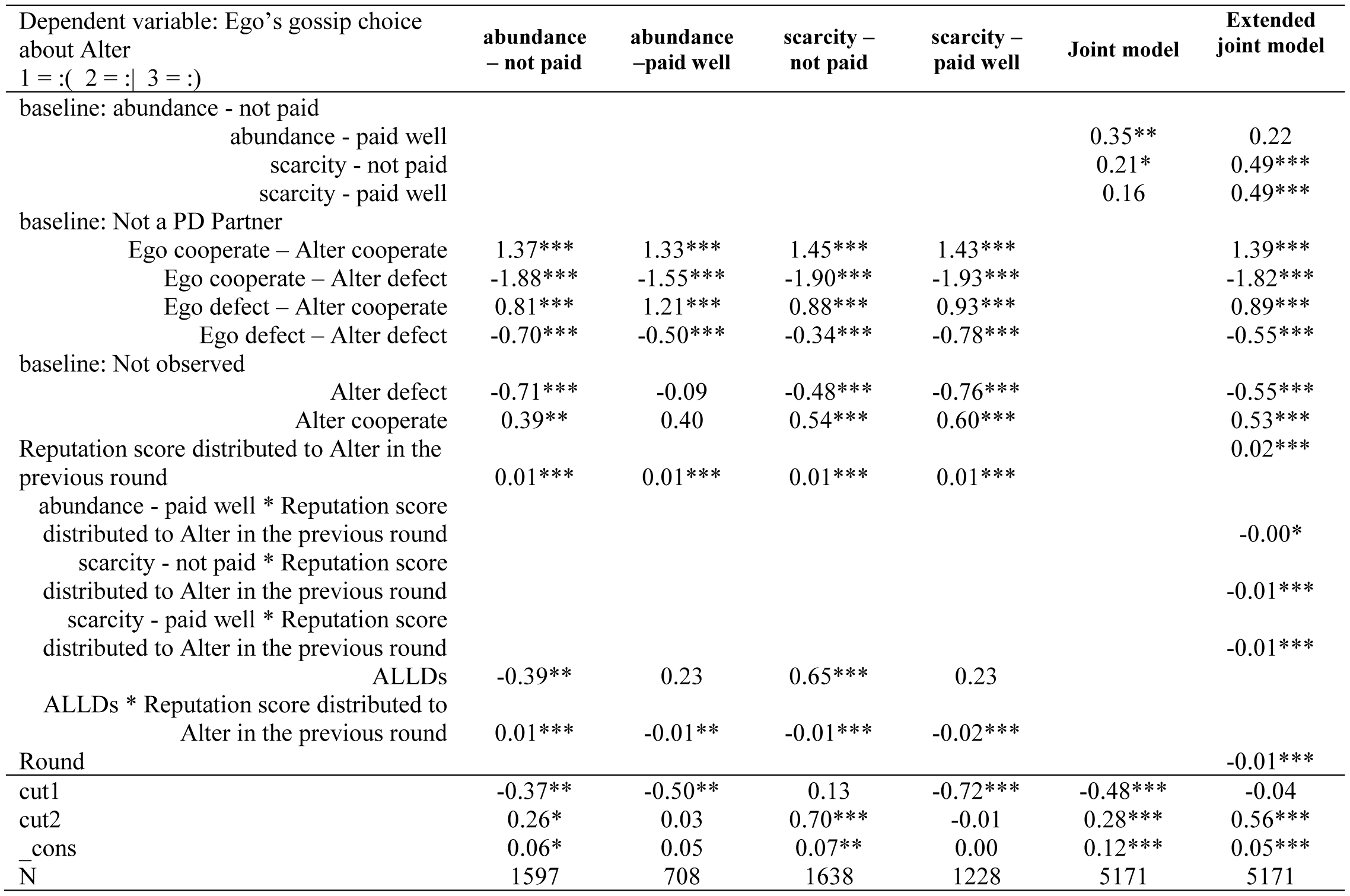
Random-effects ordered logistic models.

**Figure A1, A2.**
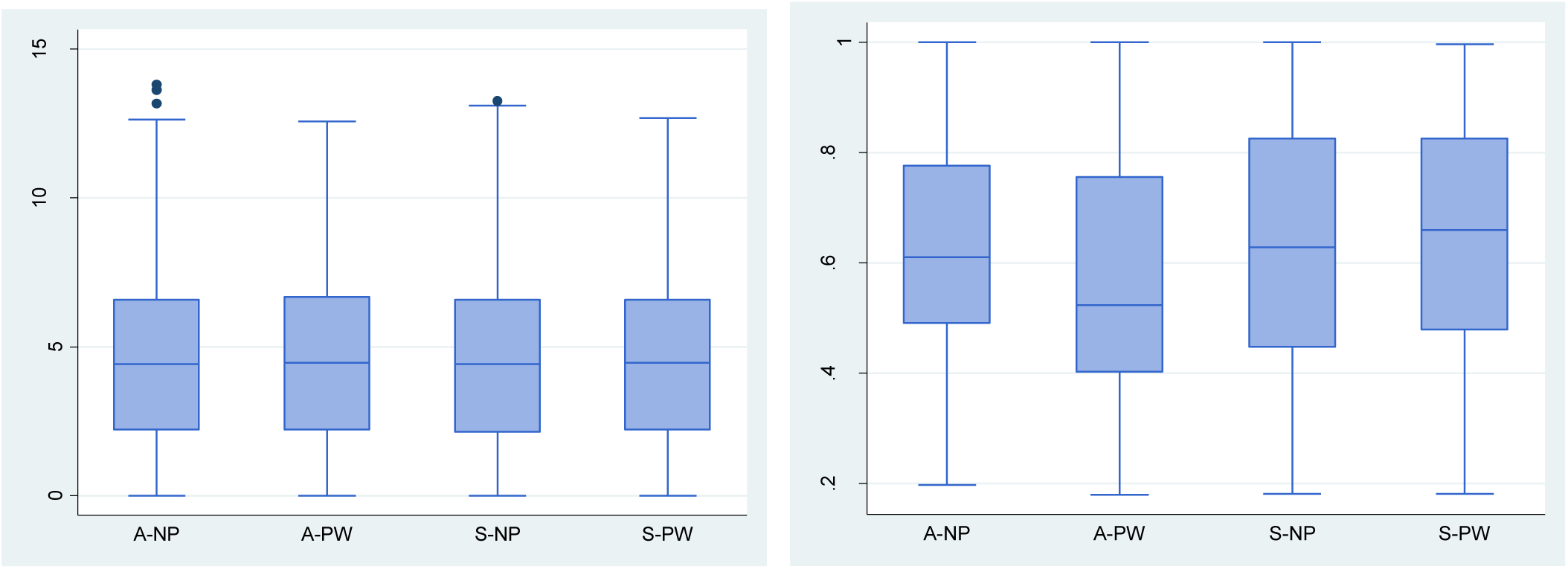

## Appendix II. Instructions of the experiment

Welcome to the decision-making experiments organized by the Corvinus University of Budapest!

The decision-making experiments carried out by the Hungarian Academy of Sciences, Centre for Social Sciences, “Lendület” Research Center for Educational and Network Studies (RECENS), led by Károly Takács, and supported by the European Research Council (ERC CoG 648693)

Please turn off your phone or completely turn it down!

In the following, instructions will appear on your screen about the experiment. You participate in the experiment together with people in this room. Instructions are the same for all participants. You get the most important instructions on paper as well. You can use them at any time during the experiment. Please do not take these instructions with you after the experiment, leave them on the table.

VERY IMPORTANT rule is that it is STRICTLY FORBIDDEN to talk to or to signal to others! Violation of this rule may result in disqualification from this experiment.

The experiment takes about 75 minutes. Your payoffs in the experiment will be paid at the end of the experiment.

The amount of your payoff depends on your own choices and the decisions of others. The precise calculation of your payment will be described later in more detail.

By pressing the \ “Next \” button, you agree that your answers will be exclusively and anonymously used for scientific research purposes.

Thank you in advance for your participation!

If you’re ready, press the \ “Next \” button! Have fun and good luck!

### INSTRUCTIONS

The experiment will consist of several rounds of decision-making. In each round, you will be paired with two other participants. Participants of the experiment will be identified with numbers ranging from 1 to 20.

It is important that you do not play with the same participants in each round! Both of your pair determines by a random number generator. The ID of your current pair will be displayed on your screen. The IDs will be kept confidential, neither during the experiment nor at the end of it we will not reveal which identifier participants belonged to,

Each round is important for your final payoff!

1 round in the first phase and 5 rounds in the second phase will be selected using a random number generator. The average winnings in these rounds will be your final payoff. We add everyone 1000 forints as a bonus. Payoffs will be rounded up to HUF 100.

Each pair faces with the following decision-making options. Two options are provided: the two options are indicated with L and R. The amount that you win this round, does not depend only on your decision, but also on your partner’s decision. It is detailed in the following what payoff can be expected:

If both of you choose L: HUF 1500

If you choose L and your partner select R: HUF 2500 If you choose R and your partner select L: HUF 0

If both of you choose R: HUF 500 If you run out of time: HUF 0

It is important that all the information that you receive is real. The time available to you will be projected in the upper right corner of your screen.

When you are ready, please click on the “Next” button.

### FURTHER INSTRUCTIONS

The same decision will be taken in the next rounds and the amounts that you can win are the same as previously.

In each round, you will randomly be paired with two other participants. It is therefore important that you do not play with the same participants in each round, but your pair is determined by a random number generator!

The change is that now the participants can be scored from 0 to 100 by you on the basis how reliable they are according to you A maximum of 950 points can be distributed. By default, the starting score is set to 50, as a neutral medium. (So long as you distribute more than the maximum points, you get an error message. If this will not be corrected in time, the total score will be rounded down proportionately.)

The points you receive will be taking into account in the final payoff! The payoff for that run will be adjusted by your average point.

If the point you received on average corresponding to the neutral value of 50, then your payoff will be unchanged in the current round. In comparison, a one-unit decrease/increase in your average point reduces/increases your payments by HUF 20. For instance, if all other participants give you 0 point, then your payment decreases by HUF 1000. If all other participants give you 100 point, your payment increases by HUF 1000.

In addition to the scoring it is also a new element that one pair from the previous rounds will be randomly selected, and you get acquainted with the decisions they have made in that round.

Then, you’ll be able to send a message to another randomly selected participant. The ID of the participant – to whom you can send the message – will appear on your screen. Then, optionally, you can select four participants, the four whom the message is about. There is no cost of sending a message.

The time available for your choice will display in the top right corner of the screen. Attention please! If you run out of time, it is considered that you did not want to send a message!

After all messages have been sent, messages you received will appear on your screen.

Now you can also modify the scores on the 100 points scale that you have given to others indicating how reliable they are. When you are ready, please click on the “Next” button. Have fun and good luck!

